# XRN1 couples RNA granule remodelling to global mRNA decay triggered by the activity of a viral endoribonuclease

**DOI:** 10.64898/2026.11.03.763857

**Authors:** Alistair Tweedie, Carla Moller-Levet, Benjamin Towler, Gillian Elliott

## Abstract

Cytoplasmic mRNA turnover is a fundamental determinant of gene expression, yet the spatial organisation of RNA decay remains unresolved. Although decay factors are enriched in processing (P)-bodies, whether these condensates actively mediate mRNA degradation or serve primarily as storage compartments remains controversial. Here, we exploit the alphaherpesvirus endoribonuclease, virion host shutoff (vhs), to synchronously trigger rapid, global cytoplasmic mRNA accompanied by nuclear relocalisation of the poly(A)-binding protein PABPC1. Expression of vhs caused the dispersal of P-bodies, but when the 5′ to 3′ exonuclease XRN1 was depleted, PABPC1 remained in the cytoplasm, and P-bodies stayed intact, associating with previously undetected G3BP1-dependent RNA granules. Similarly, catalytically inactive vhs failed to disperse P-bodies but instead induced RNA granules that also contained vhs itself and recruited decay factors in an XRN1-dependent manner. Translational shutoff by this endoribonuclease did not require RNA cleavage, G3BP1 condensation or XRN1 activity supporting a sequential model in which translational arrest precedes endonucleolytic cleavage, promoting transient RNA granules that engage P-bodies and XRN1-dependent machinery for rapid mRNA degradation within the granules, thereby precluding the biogenesis of deleterious RNA condensates.

## Introduction

mRNA decay is a major post-transcriptional mechanism for regulating the translational environment of the cell and ensuring that proteins are produced at the correct time and in the correct amount. Within the cytoplasm, the availability of mRNA for translation is regulated by translational stalling and/or mRNA decay pathways, which are influenced by membraneless protein-RNA condensates assembled through liquid-liquid phase separation (LLPS) of RNA and RNA-binding proteins (RBPs) ^1^. Processing or P-bodies (PBs), which are present under normal conditions in many cell types, are composed of RNA decay enzymes together with deadenylated mRNA ^2^. By contrast, stress granules (SGs) are not present in normal conditions but are induced by the integrated stress response (ISR) pathway or global translational arrest ^3^. SGs are made up of polyadenylated mRNAs that have failed to initiate translation, 40S ribosomal subunits and several key proteins including G3BP1 and TIA1 which make up a relatively stable internal core, and a less stable shell formed by weaker protein-protein interactions ^4,5^. PBs are frequently found docked on SGs during stress ^6^, a process that is proposed to be specific rather than the result of random collisions ^7^, and there is evidence for some components being found in both PBs and SGs ^6,8^. However, despite major advances in mRNA decay studies, it is still debated if PBs are storage sites for inactive mRNA decay enzymes or sites of decay itself, while the functional relationship between SGs and PBs remains uncertain ^6,7^.

The study of mRNA decay pathways is limited by the lack of suitable systems for inducing global mRNA degradation and often involves shutoff of transcription to track decay of mRNAs in the absence of *de novo* synthesis or the study of specific reporter transcripts ^9–12^. One elegant system that has been employed to investigate the cell biology of global mRNA decay involves the induction of the cellular antiviral endoribonuclease RNase-L which leads to widespread mRNA turnover and translational repression ^13,14^. One unusual feature of RNase-L activity is the assembly of novel RNA granules termed RLBs (RNase-L bodies) that are proposed to be distinct from SGs ^13^. However, a major limitation of this system is the requirement for RNase-L to be activated by double-stranded RNA, a canonical pathogen-associated molecular pattern (PAMP) which initiates intersecting integrated stress and interferon pathways, including RIGI/MDA5 and PKR, making outcomes specific to RNase-L-induced mRNA decay difficult to dissect ^15,16^. Moreover, RNase-L degradation is not specific for mRNA but rather for ssRNA, explaining its ability to degrade ribosomal as well as mRNA ^17,18^, a feature which further contributes to translational shutoff.

A wide range of viruses express endoribonucleases to help regulate the mRNA environment in the infected cell, favouring virus over host translation and preventing cellular responses to dsRNA ^19,20^. However, unlike RNase-L, these viral enzymes function when expressed in isolation and without the need for inducible activation. Because they hijack existing cellular processes to instigate rapid global RNA decay, they are excellent tools to reveal cellular mechanisms for the control and buffering of transcription, and the dynamics of RBPs and biomolecular condensates ^21,22^. Alphaherpesviruses express an endoribonuclease exemplified by the virion host shutoff protein from HSV1 (vhs), which targets host and viral mRNA for degradation ^23,24^. Under conditions of cytoplasmic depletion of mRNA, RBPs, including poly(A) tail-binding PABPC1, translocate to the nucleus where poly(A)+ RNA accumulates, further reducing translation ^25,26^. This effect is similar, but more complete than that seen during RNase-L activation where PABP localises to RNA granules in the cytoplasm as well as relocalising to the nucleus ^14,27^. Early studies carried out predominantly *in vitro* indicated that vhs was targeted to mRNA through binding to the translation initiation complex, causing mRNA cleavage near the 5’ end of the mRNA ^28–31^. By analogy with the gammaherpesvirus SOX protein, which cleaves mRNAs throughout the body of the transcript rather than near the 5’ end ^22^, it is hypothesized that the cellular 5’ to 3’ exonuclease XRN1 degrades the remaining uncapped mRNA ^32,33^, and mRNA fragments have been shown to persist in XRN1-depleted cells ^32^. Nonetheless, whether the vhs endoribonuclease specifically recruits XRN1 and other decay factors to accelerate decay, analogous to the recruitment of XRN1 by the cellular mRNA decapping complex ^34^ is unknown and the vhs interactome has not been defined. Likewise, the full extent of vhs activity on the cellular transcriptome has not yet been mapped.

Here, we have exploited vhs to trigger global mRNA decay and investigate its spatial organisation. We show that vhs-induced degradation of the cytoplasmic transcriptome results in the dispersal of PBs, a process that is dependent on XRN1. In the absence of XRN1, PBs are instead found associated with G3BP1-containing RNA granules that assemble only transiently when XRN1 is present. RNA granule-PB complexes were also found in cells expressing catalytically dead vhs, but in this case vhs and PB proteins were also found within the granules themselves. Hence, we propose a model of mRNA decay which requires the redirection of RNA decay enzymes away from their storage site in PBs to RNA granules containing vhscleaved RNA-protein complexes, and which only accumulate through G3BP1 condensation when either XRN1 or vhs itself are defective.

## Results

### Induction of global mRNA decay by expression of a viral endoribonuclease

We have recently shown that the vhs homologue from BoHV1 is the most active of the vhs endoribonucleases ^35^. Alphafold modelling ^36^ of vhs from HSV1 (vhsH) and BoHV1 (vhsB) revealed similar structures of the two enzymes with the N-termini of the proteins embedded in the centre of the structure (Figure 1-supplemental Figure 1). However, unlike vhs from HSV1, BoHV1 vhs has a C-terminal extension, which made it an attractive option for a GFP fusion facilitating downstream applications, without interfering with the overall protein structure or activity, a prediction supported by Alphafold modelling of the two vhs homologues C-terminally tagged with GFP (Figure 1-supplemental Figure 1). As predicted, fusion of GFP to the C-terminus of BoHV1 (vhsBGFP) vhs had no detrimental effect on vhs activity when expressed ectopically in HeLa cells, as shown by its ability to significantly reduce the expression of Gaussia luciferase (GLuc) and firefly luciferase reporters (Figure 1-supplemental Figure 1). By contrast, fusion of GFP to the C-terminus of vhs from HSV1 (vhsHGFP) greatly reduced its activity in these assays compared to untagged vhs (Figure 1-supplemental Figure 1). Moreover, BoHV1 vhs fused to GFP also induced GLuc mRNA depletion (Figure S1E) and nuclear localisation of PABPC1 (Figure 1-supplemental Figure 1) confirming that the GFP-tagged protein retained the previously identified activities of the BoHV1 and HSV1 vhs enzymes ^35^.

In line with this depletion of GLuc mRNA by BoHV1 vhs fused to GFP (hereinafter referred to simply as vhs and vhsGFP), single cell mRNA FISH of the GLuc transcript indicated that the GLuc mRNA was depleted specifically from the cytoplasm of both vhs and vhsGFP-expressing cells (Figure 1A). To confirm that this reduction in GLuc mRNA was due primarily to mRNA turnover rather than reduced transcription, nascent mRNA in HeLa cells expressing GLuc and vhsGFP was metabolically labelled with 5-Ethynyluridine (EU) at 16 hours. While the total GLuc mRNA level was significantly reduced in the presence of vhsGFP, the level of newly synthesized GLuc mRNA was unchanged (Figure 1B, total vs nascent). In this assay, treatment of cells with Actinomycin D for 1 hr significantly reduced the level of nascent GLuc mRNA, confirming that the isolation of metabolically labelled RNA had been successful (Figure 1C). Furthermore, *in situ* single-cell analysis of EU labelled cells also indicated that RNA transcription showed only slight reduction in EU labelling in cells expressing GFPvhs compared to GFP (Figure 1D).

**Figure 1.**
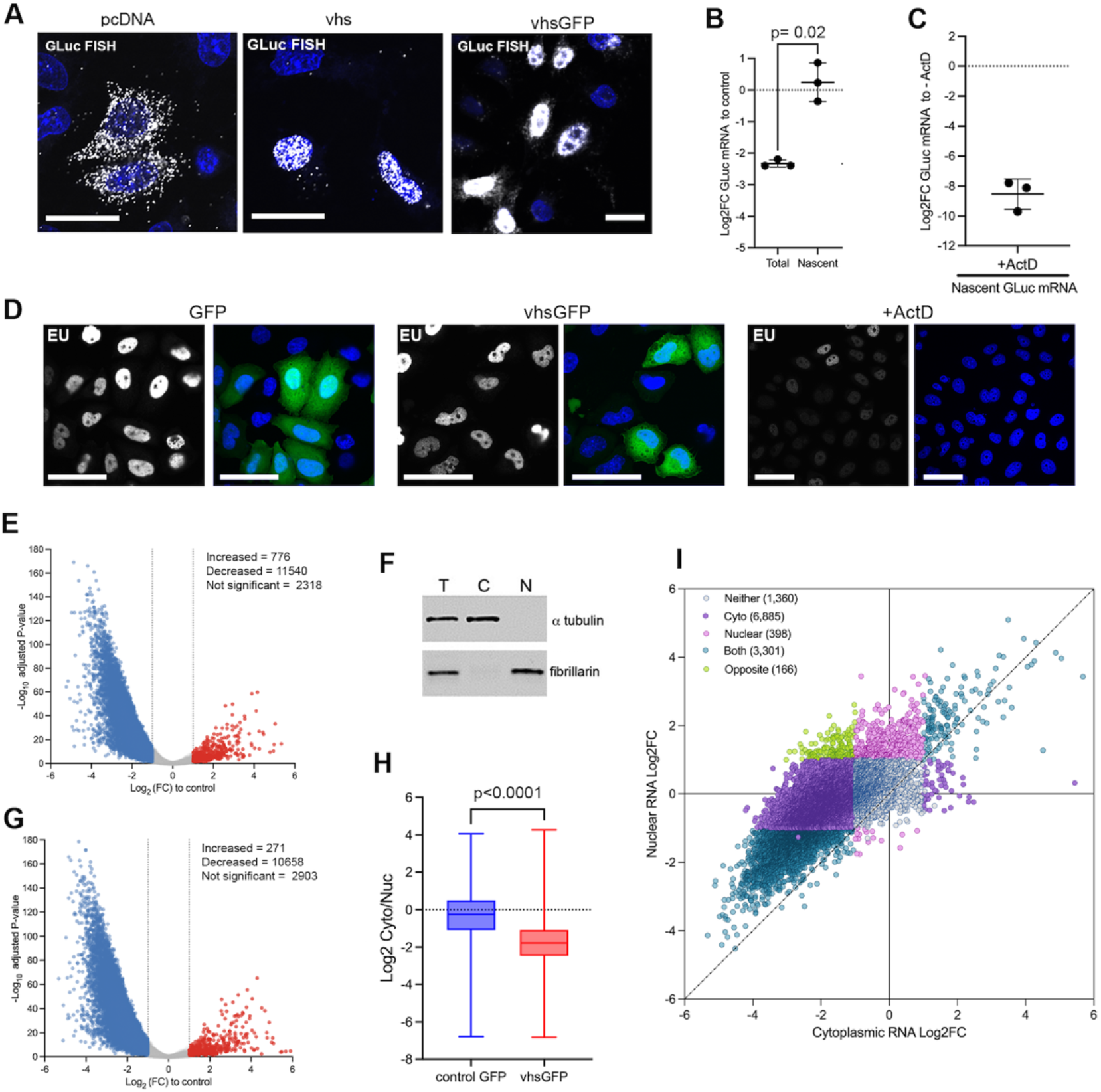
Induction of global decay of cytoplasmic mRNA by the vhs endoribonuclease. **(A)** HeLa cells transfected with plasmids expressing GLuc together with empty vector (pcDNA), plasmid expressing vhs or plasmid expressing vhsGFP were processed for mRNA FISH with probes to GLuc (white). Scale bar = 20 µm. **(B)** EU-labelled RNA from HeLa cells expressing vhsGFP and GLuc was biotinylated and purified before analysis by RT-qPCR for GLuc using 18S as reference. Two independent experiments are shown. Statistical analysis was carried out using an unpaired T-test. **(C)** HeLa cells expressing GFP and GLuc were treated with 5 µg/ml ActinomycinD (ActD) for 1 hour before EU labelling, RNA purification and GLuc amplification as in (B). Results are shown as CT values for GLuc + ActD relative to – ActD control. **(D)** HeLa cells expressing GFP or vhsGFP, or treated with 5 µg/ml ActD were labelled with EU and processed for immunofluorescence. Scale bar = 50 µm. **(E)** HeLa cells were transfected with plasmid expressing either GFP or vhsGFP, and after 16 hours GFP positive cells were sorted by FACS. Total RNA was extracted from GFP-positive cells and subjected to RNAseq analysis. The volcano plot shows the average of three independent biological repeats with significance determined as adjusted P-value <0.05. The differential expression values are available in Figure 1-Source Data 1. **(F)** Cytoplasmic and nuclear fractionation of HeLa cells were validated by Western blotting for α-tubulin (cytoplasmic marker) and fibrillarin (nuclear marker). T - total lysate; C - cytoplasmic fraction; N - nuclear fraction. **(G)** Following fractionation into cytoplasmic and nuclear fractions RNA was purified and subjected to RNAseq analysis. The volcano plot shows the average of three independent biological repeats with significance determined as adjusted P-value <0.05. The differential expression values are available in Figure 1-Source Data 1. **(H)** Box and whisper plot showing log2 cytoplasmic-to-nuclear RNA abundance ratios from RNA-seq analysis of fractionated cells expressing control GFP or vhsGFP. The dotted line at 0 denotes equal cytoplasmic and nuclear abundance. Statistical analysis was carried out using Wilcoxon test. **(I)** Scatter plot comparing the Log2FC of the nuclear transcriptome in the presence of vhsGFP and the Log2FC of the cytoplasmic transcriptome in the presence of vhsGFP. Genes are colour coded according to their behaviour in both compartments: neither, no change in cytoplasm or nucleus; cyto, DE genes in cytoplasm only; nuclear, DE genes in nucleus only; both, DE genes in cytoplasm and nucleus; opposite, DE genes that change in opposite directions. The diagonal indicates same behaviour in both compartments.

In previous work, mRNA degradation induced by vhs has generally been studied on individual reporter genes, as described above, or selected cellular genes ^37^. However, the fully active nature of the BoHV1 vhsGFP fusion protein allowed us to conduct RNAseq analysis of vhs activity to assess its overall contribution to global changes to the cellular transcriptome. HeLa cells were transfected with plasmids expressing GFP or vhsGFP and subjected to cell sorting to isolate GFP-positive cells.

RNA was then purified from whole cells and RNAseq was performed on 3 biological replicates for the GFP-positive cells in each condition, with sequence reads being mapped to the human genome, revealing that over 11,000 of 14,000 cellular genes detected were significantly reduced in response to the expression of vhs (adjusted p value < 0.05, Figure 1E and Figure 1-Source Data 1). GFP and vhsGFP expressing cells were further fractionated after cell sorting into nuclear and cytoplasmic fractions (Figure 1F) followed by RNAseq, confirming that the cytoplasmic transcriptome was depleted in vhsGFP expressing cells (adjusted p value < 0.05, Figure 1G and Figure 1-Source Data 1). In agreement with the GLuc FISH, the relative cytoplasmic-nuclear ratio of the bulk cellular transcriptome decreased significantly in vhsGFP-expressing cells compared to control GFP-expressing cells indicating reduced cytoplasmic localization (Figure 1H). Moreover, further stratification of individual genes in both compartments revealed that ~7,000 were depleted only in the cytoplasm while only ~400 increased in the nucleus, indicating that there was limited entrapment of the cellular transcriptome in the nucleus (Figure 1I). An additional 3,300 genes were depleted from both compartments which may reflect transcriptomic changes due to previously reported altered transcription as a consequence of global mRNA decay ^38–40^. This confirms that vhs expressed ectopically exhibits efficient activity on the cytoplasmic transcriptome, with both the EU labelling and the nuclear-cytoplasmic fractionation of the RNA transcriptome indicating that the outcome of vhs activity in our system appears to be predominantly post-transcriptional.

### Global mRNA decay leads to dispersal of P-bodies in an XRN1-dependent process

Given the significant depletion of the global cellular transcriptome, we next investigated the behaviour of RNA decay enzymes in the presence of vhs. Staining of HeLa cells for the major cytoplasmic 5’ to 3’ exonuclease XRN1 confirmed its localisation in PBs in untransfected and GFP transfected HeLa cells (Figure 2A). By contrast, expression of vhsGFP resulted in the effective dispersal of XRN1 from PBs (Figure 2A, compare GFP and vhsGFP). To determine if vhs affected the integrity of PBs themselves, HeLa cells expressing vhsGFP were stained for the essential PB component, the DEAD-box helicase DDX6 which colocalises with XRN1 in HeLa cells (Figure 2B, untransfected). This revealed that vhs not only dispersed XRN1 but also the entire structure of the PB (Figure 2C, vhsGFP, quantified in Figure 2D). Co-staining of cells expressing untagged vhs for PABPC1 and DDX6 (Figure 2E) further confirmed that PBs had been dispersed in the same cells where PABPC1 had been relocalised to the nucleus (quantified in Figure 2F).

**Figure 2.**
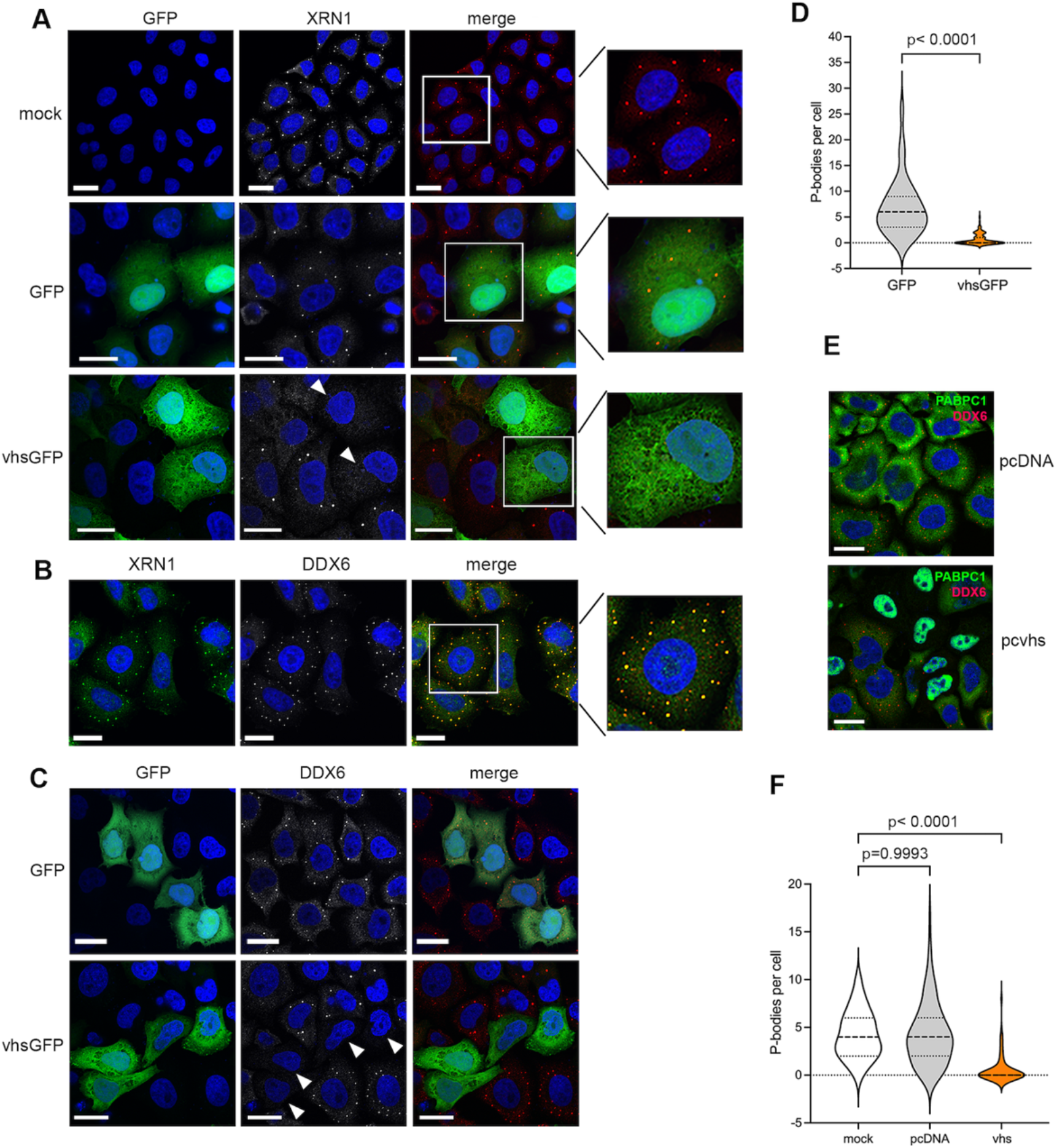
Expression of vhs causes dispersal of P-bodies. **(A)** HeLa cells left untransfected (mock) or transfected with plasmid expressing GFP or vhsGFP (green) were fixed after 16 hours and stained for XRN1 (white/red) Scale bar = 20 µm. Arrowheads, cells expressing vhsGFP. **(B)** Untransfected HeLa cells were stained with antibodies to XRN1 (green) and DDX6 (white/red). **(C)** HeLa cells transfected with plasmid expressing GFP or vhsGFP (green) were fixed after 16 hours and stained for DDX6 (white/red). Scale bar = 20 µm. Arrowheads, cells expressing vhsGFP. **(D)** P-body counts based on DDX6 staining from 100 cells for each condition across three biological repeats were conducted using ImageJ software and plotted as a violin plot showing median and quartiles. Statistical analysis was carried out using an unpaired T-test. **(E)** HeLa cells transfected with pcDNA or pcDNA expressing vhs were fixed and stained for DDX6 (green) and PABPC1 (red). Scale bar = 20 µm. **(F)** PB counts based on DDX6 staining from 100 cells for each condition across three biological repeats were conducted using ImageJ software and plotted as a violin plot showing median and quartiles. Statistical analysis was carried out using ordinary one-way ANOVA multiple comparisons, and comparison made to mock values.

To determine if XRN1 activity is required to induce nuclear relocalisation of PABPC1 in vhs expressing cells, we performed XRN1 siRNA knockdown in HeLa cells, prior to transfection with plasmid expressing vhsGFP (Figure 3A, quantified in 3B). HeLa cells depleted of XRN1 and expressing vhsGFP failed to relocalise PABPC1 to the nucleus, indicating that this outcome of vhs activity had been inhibited by the depletion of XRN1 (Figure 3C). By contrast, depletion of the closely related exonuclease XRN2 or the cellular endoribonuclease RNase-L (Figure 3A, quantified in 3B), which when activated causes PABPC1 translocation to the nucleus similar to that induced by vhs, had no effect on vhs-induced PABPC1 translocation (Figure 3C) confirming that neither of these cellular enzymes is involved in the vhs pathway. Moreover, FISH of XRN1-depleted cells expressing GLuc and vhsGFP revealed that unlike XRN1 expressing cells, where the cytoplasm was depleted of GLuc and vhsGFP transcripts (Figure 3D, sictrl), both transcripts were abundant in the cytoplasm of cells lacking XRN1 (Figure 3D, siXRN1) confirming that cytoplasmic mRNA degradation was abrogated when XRN1 was depleted. Intriguingly, puromycin labelling of cells expressing vhsGFP revealed that translational shutoff induced by vhs was not dependent on the presence of XRN1 or PABPC1 localisation to the nucleus (Figure 3E). This indicates that unlike RNase-L ^14^, translational shutoff induced by vhs is not a direct consequence of RNA degradation and that the translational shutoff and mRNA degradation properties of vhs are separable, as suggested previously^35^.

**Figure 3.**
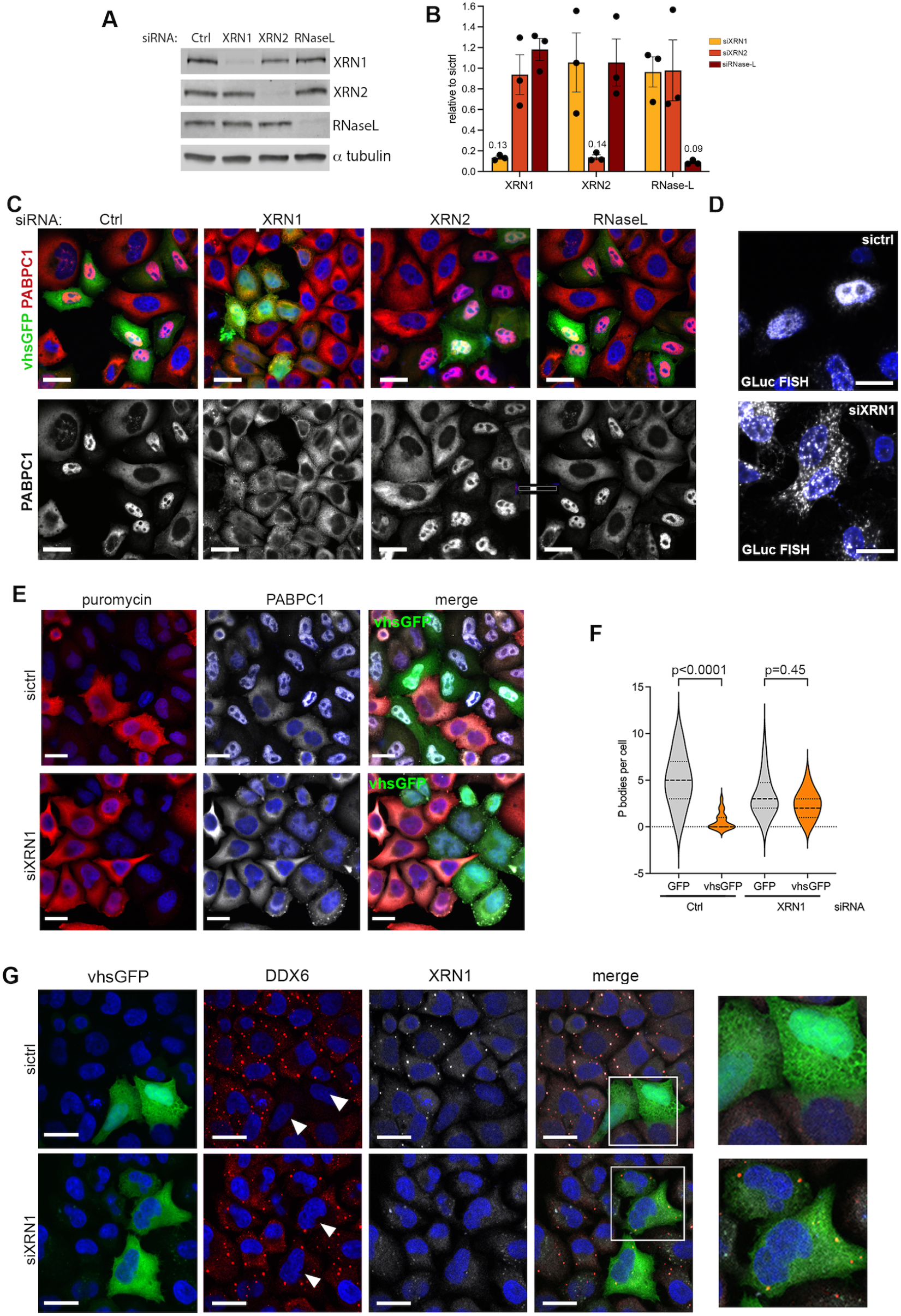
P-body dispersal by vhs is dependent on XRN1. **(A)** HeLa cells transfected with a non-targeted control siRNA (ctrl) or siRNA to XRN1, XRN2 or RNase-L were harvested after 48 h and analysed by Western blotting with antibodies as indicated. **(B)** Three biological replicates of (A) were normalised to a tubulin and quantified relative to ctrl siRNA. Error bar = SEM. **(C)** HeLa cells transfected with control or XRN1, XRN2 or RNase-L siRNAs were further transfected after 48 h with plasmid expressing vhsGFP. Cells were fixed after a further 16 hours, and stained for PABPC1 (red/white). Scale bar = 20 μm. **(D)** HeLa cells that had been transfected with siRNA to XRN1 were subsequently transfected with plasmids expressing vhsGFP and GLuc. After a further 16 hours, the cells were processed for mRNA FISH using probes against GFP (green) and GLuc (red). Scale bar = 20 µm. **(E)** HeLa cells transfected with control or XRN1 siRNAs were transfected with plasmid expressing vhsGFP and labelled with puromycin 16 h later. Cells were fixed and stained with antibodies to puromycin (red) and PABPC1 (white). Scale bar = 20 μm. **(F) & (G)** HeLa cells transfected with control or XRN1 siRNAs were further transfected after 48 h with plasmid expressing vhsGFP. Cells were fixed after a further 16 hours, and stained for DDX6 and XRN1. DDX6-positive P-body counts from 100 cells for each condition were carried out using ImageJ software and plotted as a violin plot showing median and quartiles (F). Statistical analysis was carried out using ordinary one-way ANOVA multiple comparisons and comparisons made to GFP ctrl siRNA values. Representative fields showing DDX6 (red) and XRN1 (white) are shown in (G). Scale bar = 20 μm. White arrowheads indicate cells expressing vhsGFP.

DDX6 staining of XRN1-depleted cells showed that PBs remained intact in the absence of XRN1 (Figure 3-supplemental figure 1), although their numbers were reduced from an average of 5 to an average of 3 per cell (Figure 3-supplemental figure 1). Strikingly, expression of vhsGFP failed to disperse PBs in XRN1-depleted cells (Figure 3G, vhsGFP; quantified in Figure 3F) with the small reduction in average PB number from 3 to 2 in the presence of vhsGFP potentially reflecting an incomplete depletion of XRN1 in some cells. Hence, PB dispersal by the vhs endoribonuclease is dependent on XRN1.

### RNA granules form next to P-bodies in XRN1-depleted cells expressing vhs

In normal conditions, the expression of vhs leads to the depletion of cytoplasmic polyadenylated RNA reflecting the activity of XRN1 on vhs-cleaved mRNA ^35^ and (Figure 4A, sictrl). By contrast, poly(A)+ FISH of XRN1-depleted cells expressing vhs (Figure 4A, siXRN1) or vhsGFP (Figure 4B) revealed that polyadenylated RNA was no longer depleted from the cytoplasm, but instead, multiple small poly(A)+ enriched cytoplasmic RNA foci had formed.. Staining of XRN1-depleted, vhs-expressing cells confirmed that these granules contained the SG nucleating protein G3BP1 (Figure 4C), while closer examination of non-depleted cells expressing vhs stained for G3BP1 also revealed small foci in a subset of cells where vhs had induced dispersal of DDX6 (Figure 4D); we suggest that these foci are a transient form of the G3BP1 foci seen above in XRN1-depleted cells. Additionally, co-staining of XRN1-depleted, vhs-expressing cells for PABPC1 and TIA1, both of which are known to localise to canonical SGs ^3^, revealed that vhs-induced condensates also contained both of these proteins (Figure 4E & 4F), while triple staining of non-depleted cells revealed colocalization of G3BP1, PABPC1 and TIA1 in cells where these small foci were detectable (Figure 4G). We therefore propose that these condensates are similar to nascent SGs, but rather than being triggered by translational arrest through the ISR pathway, these vhs-induced SGs have been induced through direct translational blockade by vhs. In the absence of XRN1, these bodies appear to be stabilised. While conducting this imaging, we also noted that in line with the known nuclear shuttling of TIA1 ^41^, TIA1 became more concentrated in the nuclei of cells expressing vhsGFP (Figure 4G & 4H, TIA1), suggesting that like PABPC1, the shuttling dynamics of TIA1 had been altered by the depletion of cytoplasmic mRNA.

**Figure 4.**
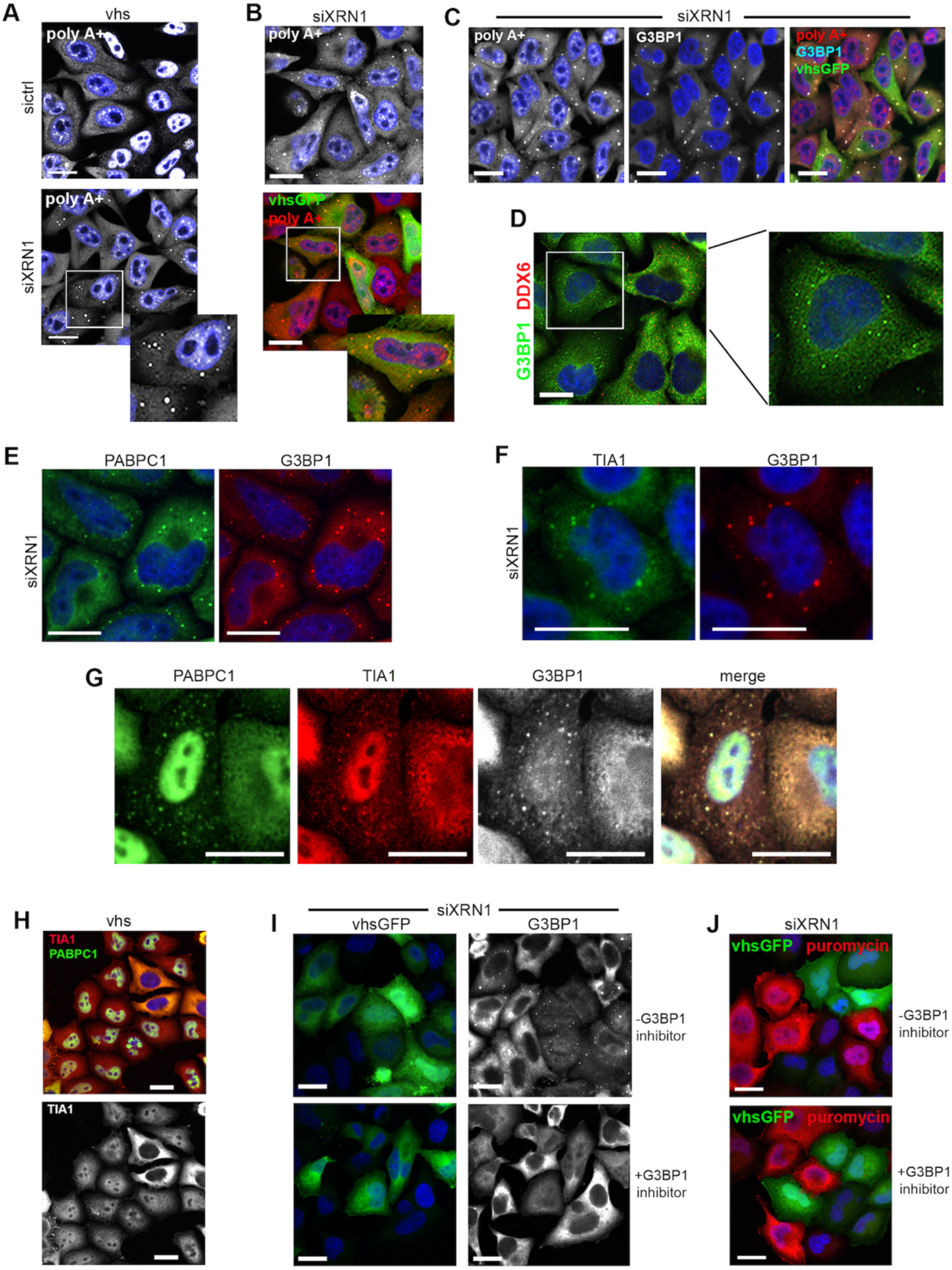
Vhs induces G3BP1-dependent RNA granules in cells lacking XRN1. **(A) & (B)** HeLa cells were transfected with control or XRN1 siRNA, and after 48 hours further transfected with plasmid expressing untagged vhs (A) or vhsGFP (B). Cells were then fixed after a further 16 hours, and processed for poly(A)+ FISH (white/red). Scale bar = 20 µm. **(C)** As for (B), but cells were also stained for G3BP1 (white/cyan). Scale bar = 20 µm. **(D)** HeLa cells expressing untagged vhs by transient transfection were stained for G3BP1 (green) and DDX6 (red). Scale bar = 20 μm. **(E) & (F)** HeLa cells that had been transfected with XRN1 siRNA were subsequently transfected with plasmid expressing vhsGFP. After a further 16 hours, the cells were fixed and stained for G3BP1 and PABPC1 (E), or G3BP1 and TIA1 (F). Scale bar = 20 μm. **(G)** HeLa cells expressing untagged vhs by transient transfection were stained for PABPC1 (green), TIA1 (red) and G3BP1 (white). Scale bar = 20 μm. **(H)** HeLa cells expressing untagged vhs by transient transfection were stained for PABPC1 (green) and TIA1 (red). Scale bar = 20 μm. **(I)** HeLa cells that had been transfected with XRN1 siRNA were subsequently transfected with plasmid expressing vhsGFP in the absence of presence of the G3BP1 inhibitor G31b (50 µm). After a further 16 hours, the cells were fixed and stained for G3BP1 and PABPC1. Scale bar = 20 µm. **(J)** HeLa cells that had been transfected with XRN1 siRNA were subsequently transfected with plasmid expressing vhsGFP in the absence of presence of the G3BP1 inhibitor G31b (50 µm). After a further 16 hours, the cells were labelled with puromycin, fixed and stained for puromycin (red). Scale bar = 20 µm.

To determine if G3BP1 condensation is required for these granules to form, we utilised the G3BP1 inhibitor G31b, a drug which maintains G3BP1 ability to interact with RNA but disrupts interactions between G3BP1 and other SG proteins that are essential for driving liquid-liquid phase separation during SG condensation ^42^. G31b was confirmed to prevent formation of canonical stress granules induced by heat shock (Figure 4-supplemental figure 1) while having no effect on non-stressed cells (Figure 4-supplemental figure 1). G31b also prevented formation of RNase-L-induced bodies in cells treated with poly(I:C) (Figure 4-supplemental figure 1); in this scenario, the localisation of PABPC1 to the nucleus induced by RNase-L mRNA cleavage was unaffected by inhibition of G3BP1 condensation. Treatment of cells expressing vhsGFP with G31b indicated that G3BP1 condensation is also required for assembly of these granules in XRN1-depleted cells (Figure 4I). Labelling with puromycin also indicated that G3BP1 condensation was not required for translational shutoff induced by vhs, confirming that translational blockade occurs upstream and is not dependent on granule formation (Figure 4J), as shown previously for canonical stress granules ^42^.

To determine the relationship between these endoribonuclease-induced granules and PBs, XRN1-depleted cells expressing vhsGFP were stained for G3BP1 and DDX6 (Figure 5A). Low magnification imaging revealed that PBs were frequently found juxtaposed to G3BP1 granules (Figure 5A, white arrowheads), suggesting that PBs had been recruited to the small vhs-induced RNA granules, or vice versa. Higher magnification imaging of individual G3BP1-PB complexes demonstrated this more clearly (Figure 5B & 5C), with representative line profile analysis for DDX6 and G3BP1 indicating that although closely abutted, these bodies did not coincide (Figure 5D). Counts of PBs and vhs-induced granules in vhsGFP expressing cells revealed that around 50% of the vhs-induced granules were found to be associated with PBs (Figure 5E). G3BP1 and DDX6 staining of XRN1-depleted cells expressing untagged vhs confirmed this phenotype, indicating that the GFP tag on vhs played no part in this localisation (Figure 5F).

**Figure 5.**
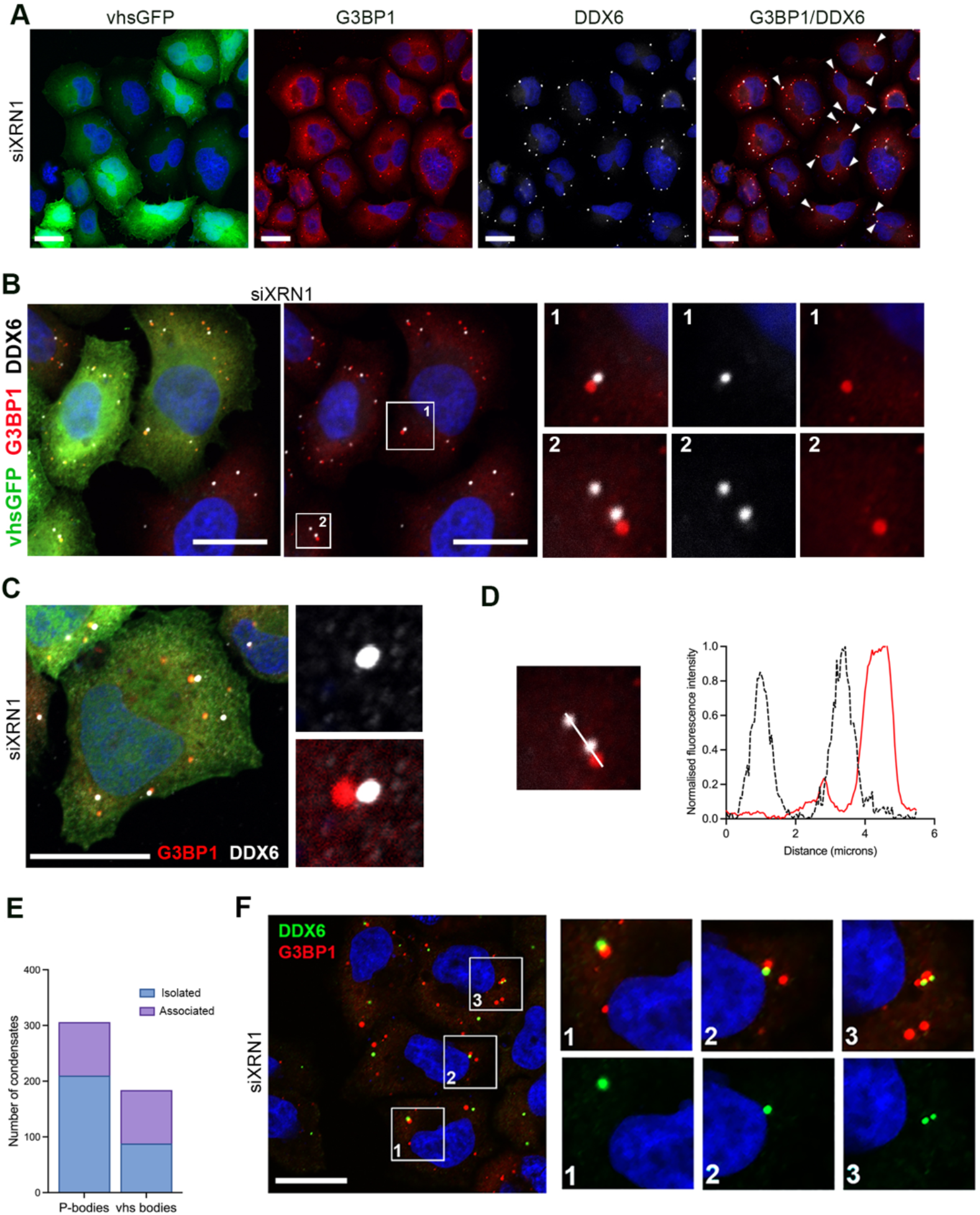
P-bodies associate with vhs-induced G3BP1 granules. **(A)** HeLa cells were transfected with XRN1 siRNA, and after 48 hours further transfected with plasmid expressing vhsGFP. Cells were fixed after a further 16 hours and stained for DDX6 (white) and G3BP1 (red). Scale bar = 20 µm. White arrowheads point to P-bodies juxtaposed to G3BP1 bodies. (B) As for (A) but higher magnification. (C) As for (B). (D) Line profile analysis of DDX6 (white) and G3BP1 (red) from panel 2 in (B). (E) P-bodies and vhs-induced granules (vhs bodies) were counted in individual cells and scored for being isolated or associated, n=50. (F) As for (A) but XRN1 depleted cells were transfected with plasmid expressing untagged vhs. Cells were fixed after a further 16 hours and stained for DDX6 (green) and G3BP1 (red). Scale bar = 20 µm.

### Catalytically dead endoribonuclease causes accumulation of mRNA decay factors in G3BP1-containing granules

Several residues thought to be important for cleavage of RNA by vhs have previously been identified ^43^, and are highly conserved across the vhs homologues (Figure 6-supplemental figure 1). Modelling their location using Alphafold revealed them to be embedded in the central region of the vhs protein structure, with Mg^2+^ binding site 1 (dark blue in Figure 6A, with accessory threonine residue thought to be important for activity in light blue) and Mg^2+^ binding site 2 (green in Figure 6A) located close to each other and the N-terminus of the protein (maroon in Figure 6A). In the case of the T211A mutant (light blue in Figure 6A), this residue has been proposed to be important for activity and hydrogen bonds with D34^43^. Muta-genesis of these residues (D34N, D83N, D213N, D215N and T211A) was performed in the vhsGFP background to generate vhs^*dead*^ mutants, and expression confirmed by Western blotting of cell extracts for GFP (Figure 6B). As shown before ^35,44^, vhs was expressed at a low level, a feature we have hypothesized is due to it inducing the degradation of its own mRNA ^37^. Of note, for vhs variants with point mutations of D34N, D83N, D213N or D215N, expression of the vhsGFP fusion proteins was partly or wholly recovered to the level of GFP itself (Figure 6B) and this was reflected in the relative transcript levels for each of the variants (Figure 6C, green), supporting the hypothesis that vhs regulates its own expression. The relative effect of these mutations on the decrease of the GLuc reporter transcript when expressed in the presence of vhs (Figure 6C, red), confirmed that these residues are important for mRNA decay.

**Figure 6.**
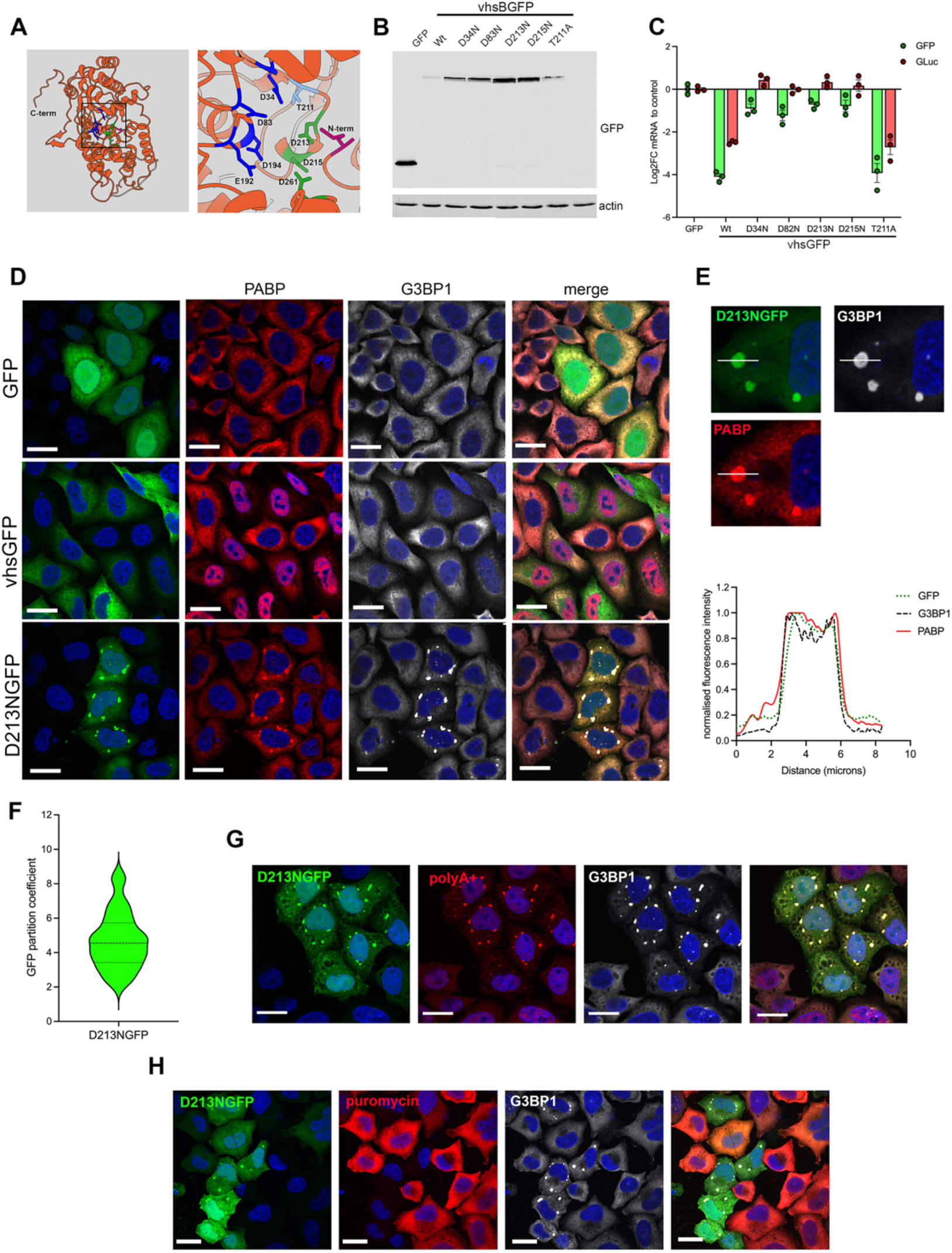
Catalytically dead vhs induces G3BP1 condensates. **(A)** Alphafold modelling of vhs showing that the catalytic site is embedded in the centre of the protein structure. Important residues for catalysis are highlighted according to ^43^: blue – Mg binding site 1; green – Mg binding site 2; light blue - threonine residue thought to be important for activity and hydrogen bonds with D34 in Mg binding site 1; maroon – N-terminal methionine. **(B)** HeLa cells transfected with plasmids expressing vhsGFP variants or GFP alone as indicated were subjected to SDS-PAGE and Western blotting with antibodies to GFP and b-actin. **(C)** Plasmids expressing GFP-tagged active site mutants of vhs were co-transfected with the GLuc reporter plasmid as indicated and GFP and GLuc expression measured by RT-qPCR using 18S as reference (n=3). **(D)** HeLa cells expressing GFP-tagged D213N variant of vhs (green) were fixed and stained 16 hours later for PABPC1 (red) and G3BP1 (white). Scale bar = 20 µm. **(E)** Line profile analysis of normalized fluorescence intensity for GFP, G3BP1, and PABP across the indicated granule in left hand panels. Intensities were normalized to the maximum value in each channel. **(F)** Violin plot of partition coefficient measurements for D213NGFP as measured using ImageJ software. The dashed line indicates the median and dotted lines indicate the interquartile range. n=80. **(G)** HeLa cells were transfected with plasmid expressing D213NGFP then fixed after 16 hours and processed for poly(A)+ FISH (red) and stained with antibody to G3BP1 (white). Scale bar = 20 µm. **(H)** HeLa cells transfected with plasmid expressing D213NGFP were labelled with puromycin 16 h later. Cells were fixed and stained with antibodies to puromycin (red) and G3BP1 (white). Scale bar = 20 μm.

When expressed in HeLa cells, the D34N, D83N, D213N and D215N mutants were unable to relocalise PABPC1 to the nucleus, indicative of a failure to induce mRNA degradation (exemplified by D213N in Figure 6D; Figure 6-supplemental figure 2), while T211A maintained this function (Figure 6-supplemental figure 2). D34N, D83N, D213N and D215N induced large cytoplasmic granules exemplified by D213N in Figure 6D; Figure 6-supplemental figure 2) which contained not only PABPC1 and G3BP1 but also TIA1 (Figure 6-supplemental figure 3). Additionally, vhsGFP itself was concentrated in these granules (Figure 6D & line profile analysis shown in Figure 6E), with an average partition coefficient of around five (Figure 6F), while poly(A)+ FISH confirmed that they were greatly enriched in polyadenylated mRNA (Figure 6G). Puromycin labelling of cells further showed that despite not causing mRNA degradation, catalytically dead mutant D213NGFP maintained the ability to shut off translation in a manner similar to Wt vhsGFP (Figure 6H). This suggests that catalytically dead variants of vhs retain the ability to interact with mRNA-RBP complexes resulting in vhs-induced translational shutoff and the assembly of larger SGs.

Having shown that active vhs induces dispersal of PBs, we next investigated what happened to PBs in cells expressing vhs^*dead*^. Unlike active vhs, expression of D213N (or the other vhs^*dead*^ mutants) did not disperse PBs. Instead, these were frequently found juxtaposed to the D213N-induced RNA granules (Figure 7A), in a similar manner to XRN1-depleted cells expressing Wt vhs (Figure 5). However, in contrast to Wt vhs, expression of D213N led to the accumulation of DDX6 within these G3BP1 granules (Figure 7A). When PBs were associated with these G3BP1 granules, DDX6 was found in three different patterns: a single PB associated with DDX6-positive granules (Figure 7A, top row), multiple PBs associated with DDX6-positive granules (Figure 7A, middle row) or DDX6-postive granules without any obvious associated PBs (Figure 7A, bottom row), as shown by line profile analysis of representative DDX6-positive granules (Figure 7B). DDX6 had a partition coefficient of around 5 in cells expressing catalytically inactive vhs, either untagged or GFP-tagged (Figure 7C). Moreover, XRN1 itself was also shown to be recruited to these vhs^*dead*^-induced granules (Figure 7D) suggesting that the mRNA decay machinery had been specifically recruited. By contrast, when XRN1 was depleted from cells expressing vhs^*dead*^, PBs were still found associated with the vhs^*dead*^-induced granules but there was no detectable DDX6 within them (Figure 7E & 7F) as confirmed by line profile analysis of representative granules (Figure 7GF). These conditions phenotyped poly (I:C) treated cells where RNaseL-induced RLBs were found juxtaposed to PBs with no evidence for redirection of XRN1 or DDX6 into RLBs (Figure 7-supplemental Figure 1). As RNase-L does not activate XRN1 degradation of mRNA ^50, 51^, RLBs would not be expected to recruit the mRNA decay machinery. Taken together, these data show that XRN1 is essential for the redirection of the decay machinery to nascent vhs-induced granules, resulting in rapid mRNA decay and PB dispersal.

**Figure 7.**
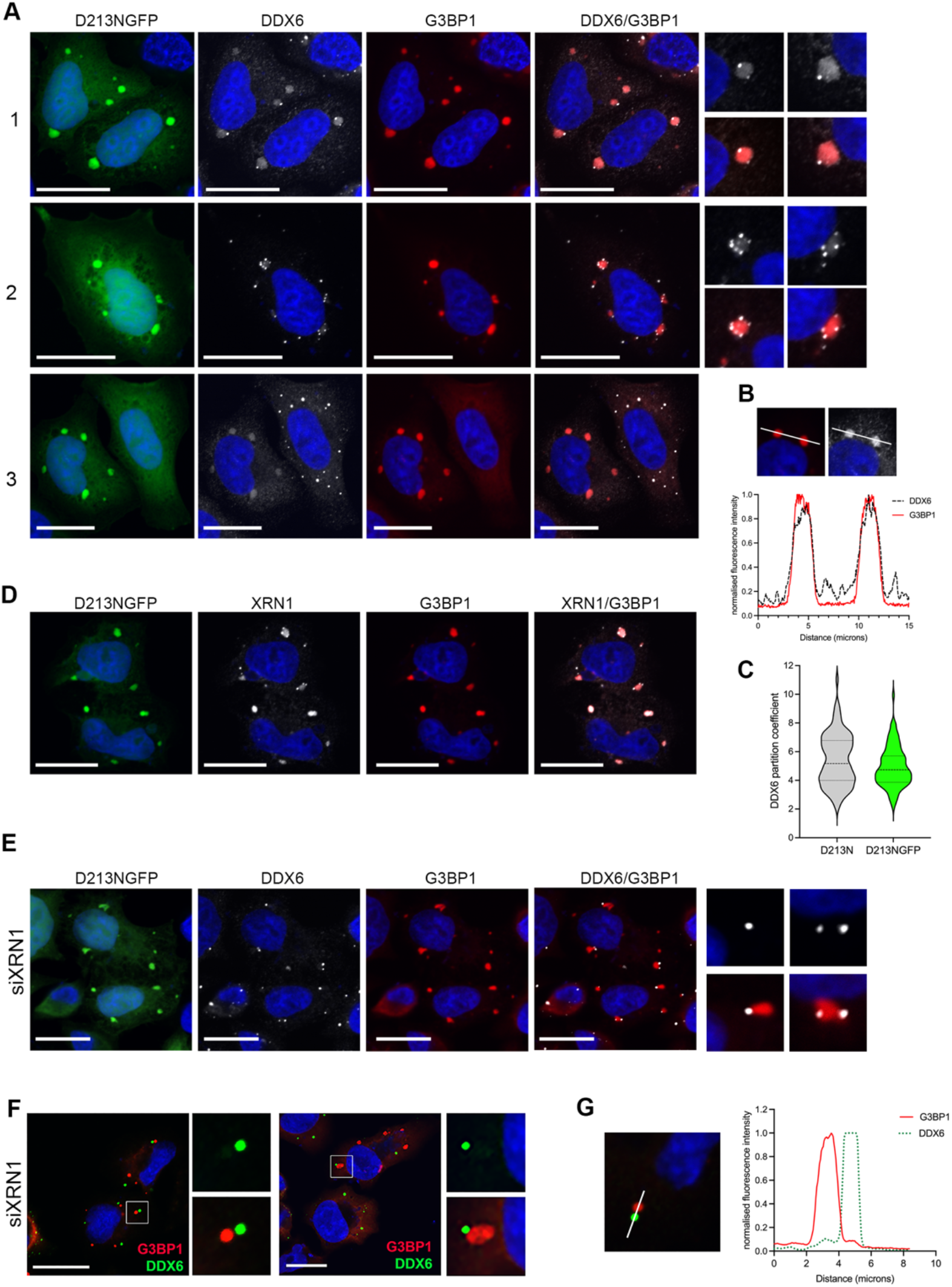
P-body components concentrate in vhs-induced RNA condensates in an XRN1 dependent process. **(A)** HeLa cells grown on co-verslips were transfected with plasmid expressing vhsGFP D213N and fixed after 16 hours. Cells were labelled with antibodies to DDX6 (white) and G3BP1 (red). Scale bar = 20 µm. (**B)** Line profile analysis of normalized fluorescence intensity for DDX6 (black) and G3BP1 (red) across the indicated granules in top panel. Intensities were normalized to the maximum value in each channel. **(C)** Violin plot of partition coefficient measurements for DDX6 in granules formed after expression of untagged D213N and D213NGFP. The dashed line indicates the median and dotted lines indicate the interquartile range. n=90. **(D)** As for (A), but cells were stained for XRN1 (white) instead of DDX6. **(E)** HeLa cells that had been transfected 48 hours earlier with siRNA to XRN1 were transfected with plasmid expressing D213NGFP, fixed after 16 h and stained for G3BP1 (red) and DDX6 (white). Scale bar = 20 µm. **(F)**. As for (G), but cells were transfected with plasmid expressing untagged D213N and stained for G3BP1 (red) and DDX6 (white). Scale bar = 20 µm. **(G)** Line profile analysis of normalized fluorescence intensity for DDX6 (green) and G3BP1 (red) across the indicated granules in left hand panel. Intensities were normalized to the maximum value in each channel.

## Discussion

Given its global activity and exploitation of the cellular mRNA decay machinery, the viral endoribonuclease vhs provides an excellent model system for delineating the subcellular compartmentalisation of mRNA decay that would otherwise be intractable at the individual mRNA level. It was first proposed that mRNA decay occurred in PBs, because cellular RNA decay factors accumulate there ^45^. Later work showing that RNA decay was not affected by mutants unable to form PBs suggested a role for them in storage rather than decay ^11^, but recently, this debate has been reopened with new evidence showing rapid mRNA decay inside PBs ^10^. Here we show that synchronous mRNA decay triggered by the endonucleolytic activity of vhs reveals a dynamic communication between P-bodies and SG-like condensates that is controlled by XRN1 and ultimately prevents the accumulation of aberrant stress-associated RNA granules.

Our data leads us to propose a sequential pathway for global mRNA decay that involves recruitment of RNA decay enzymes into condensates containing cleaved poly(A)+ RNA, potentially directly from associating PBs, resulting in the dispersal of both types of condensate. In this model, vhs binds to the 5’ end of mRNAs; as vhs is not an RBP, it is likely to be brought into proximity with the RNA by interacting with components of the 5’ cap binding complex, such as the previously proposed binding partners EIF4H or EIF4A ^46^. In our model, this results in translational blockade, leading to the formation of nascent RNA granules through G3BP1 condensation, followed by PB recruitment and XRN1-dependent localisation of RNA decay enzymes into these granules. If mRNAs in these granules have been cleaved, then rapid XRN1-mediated degradation occurs and the granules and PBs disperse rapidly. However, if XRN1 is not present, then the granules are stabilised but do not grow larger because vhs continues to cleave mRNA and G3BP1 fails to condense a larger network around these RNA fragments. Additionally, if the granules contain catalytically dead endoribonuclease, then the mRNAs are not cleaved and the granules expand further to resemble canonical SGs while still recruiting RNA decay enzymes. In line with the previously proposed stepwise formation of SGs ^8^, this would suggest that vhs-induced RNA decay within small granules actively prevents the biogenesis of larger RNA granules formed through initial translational blockade. As there are only a handful of mRNAs left untouched by vhs when it is expressed in isolation, translational shutoff would therefore be maintained in these cells through mRNA degradation, as evidenced by our puromycin labelling studies shown here (Figure 3E) and elsewhere ^47^.

Translational arrest is generally induced by one of two mechanisms in the cell: activation of the ISR pathway leading to phosphorylation of the translation initiation factor eIF2α and the assembly of canonical SGs; or direct targeting of the eIF4F complex involved in translation initiation and the assembly of non-canonical SGs ^3^. In the case of vhs expression, translational shutoff did not require G3BP1 condensation, XRN1 activity or RNA cleavage by vhs, placing shutoff at the start of the vhs mRNA decay pathway, in line with the non-canonical SG pathway. Moreover, vhs-induced granules are similar in size and content to SG cores that form in the initial stage of SG assembly ^5^ suggesting that they may represent the initial SG seed but are limited in size by mRNA fragmentation preventing extensive G3BP1 nucleation events. However, if vhs itself is unable to cleave the mRNA, SG nucleation by G3BP1 continues and the granules increase in size. By contrast, work on the cellular endoribonuclease RNase-L which responds to the presence of aberrant RNA has shown that catalytically dead RNase-L does not form RLBs, suggesting that unlike vhs, both translational arrest and RLB assembly are a consequence of mRNA cleavage ^13^.

Detailed proteomic analysis of SGs and PBs has shown that they share some common components ^53^, but how these proteins are recruited into each biocondensate is still unclear. Protein transfer from PBs into SGs has been proposed before ^6^, and mRNA decay in SGs has recently been shown to be dependent on XRN1 ^48^, but to our knowledge our data presented here is the first indication that RNA decay enzymes such as DDX6 and XRN1 are recruited into SGs in an XRN1-dependent process. By contrast, others have shown that DDX6 is not present in SGs that have formed as a consequence of stress caused by exposure to arsenite, which induces the ISR pathway ^49^. The formation of vhs-induced RNA granules also has parallels with the RLBs formed when RNase-L has been activated by dsRNA; small G3BP1-containing granules form, PABPC1 relocates to the nucleus and PBs are found associated with these RLBs ^13,14^. Moreover, unlike the situation when active vhs is expressed, PBs remain intact and decay enzymes are not redirected to RLBs when PBs and RLBs interact (Figure 7-supplemental Figure 1) ^13^. In line with this and unlike vhs, RNase-L activity does not recruit the 5’ to 3’ cellular exonuclease XRN1 for rapid mRNA degradation. Although RNase-L cuts at multiple places through mRNAs, it leaves 5′-OH ends rather than the 5’-monophosphates required by XRN1 such that any subsequent degradation by XRN1 is significantly delayed or does not occur ^50,51^. Hence, RNase-L induced RNA fragments accumulate in RLBs leading to subsequent RIGI recognition and further activation of the IFN pathway ^52^. While important for sensing of virus infection, it would be highly detrimental for an endoribonuclease to similarly activate IFN in this way in the normal context of cellular mRNA decay, and hence the ability to recruit XRN1 and other mRNA decay factors would prevent deleterious activation of IFN. It has recently been shown that XRN1 depletion results in the inhibition of mRNA decapping by disrupting the activity of the scaffold protein EDC4, a protein which enhances mRNA decapping by the enzyme DCP2, resulting in translationally repressed mRNA ^54^. The similarity of this phenotype with the phenotype of XRN1-depleted, vhs-expressing cells shown here indicates that the global mRNA depletion induced by vhs closely reflects the basal decay mechanism seen during normal mRNA decapping and turnover in the cellular environment, further emphasizing the suitability of vhs as a model for mRNA decay. The use of catalytically dead vhs has allowed us to “freeze” the dynamics of the PB-vhs granule interaction, thereby providing a future means of delineating the subcellular location and protein-protein interactions involved in mRNA decay, and new information on the as yet undefined pathway for PB disassembly ^55^. Because much of the previous work on vhs has been conducted *in vitro*, the outstanding questions on the molecular detail of this process – the binding partners that allow it to interact with mRNA, the mRNA cleavage recognition site, how many times cleavage occurs within a single transcript, and how it recruits XRN1 – should now be revisited to enhance the mechanistic understanding of mRNA decay. Finally, although out of the scope of this current study, the relative susceptibility of cellular mRNAs to endoribonuclease activity as indicated by our transcriptome data (Fig 1I and Figure 1-Source Data 1) will reveal additional detail on the relative metabolism and fate of individual mRNAs, opening up the potential for a systems approach to understanding mRNA decay and the normal turnover of individual cellular mRNAs.

## Materials and Methods

### Cells and plasmids

HeLa cells were cultured in DMEM (Invitrogen # 11504496) supplemented with 10% foetal bovine serum (Fisher Scientific # 11550356). Plasmid pEGFPN1 expressing GFP under the control of the hCMV immediate-early promoter was obtained from Clontech. The construction of plasmids expressing vhs from HSV1 strain17 or BoHV1 strain P8-2 as GFP-tagged and untagged versions have been described previously ^35^. Vhs^*dead*^ mutations in BoHV1 vhs were introduced by site-directed mutagenesis (New England Biosciences kit # C2987H) and confirmed by sequencing. Plasmids were transfected into HeLa cells using the Lipofectamine 2000 reagent (Invitrogen # 10696153), and cells were routinely harvested 16 h later.

## Antibodies

Commercial antibodies used in this study were mouse α-tubulin (Sigma # T1568), mouse GFP (Proteintech # 66002-1), mouse XRN1 (Santa Cruz # sc-165984), rabbit DDX6 (Bio-Techne # NB200-191), mouse PABPC1 (Santa Cruz # sc-32318), rabbit PABPC1 (Abcam # ab21060), mouse G3BP1 (Abcam # ab56574), rabbit fibrillarin (Abcam # ab5821), goat TIA1 (Santa Cruz # sc-1751), mouse RNaseL (Santa Cruz # sc-74405), mouse XRN2 (Santa Cruz # SC-365258) and mouse puromycin (EMD Millipore # MABE343).

### SDS-PAGE and Western blotting

Protein samples were analysed by SDS-polyacrylamide gel electrophoresis and transferred to nitrocellulose membrane for Western blot analysis. IRDye secondary antibodies were obtained from LI-COR and Western blots were imaged on a LI-COR Odyssey Imaging system.

### Drugs

The G3BP1 inhibitor FAZ-3780 (Cambridge Biosciences # Hy-162289) was used at 50 µM. Actinomycin D (Sigma) was used at 5 µM to inhibit RNA polymerase I and II transcription and 50 nM to inhibit RNA polymerase I transcription alone.

### Immunofluorescence

Cells for immunofluorescence were grown on coverslips and fixed with 4% paraformaldehyde (PFA) in PBS for 20 min at room temperature, followed by permeabilisation with 0.5% Triton-X100 (Merck) for 10 min. Fixed cells were blocked by incubation in PBS with 10% newborn calf serum for 20 min, before the addition of primary antibody in PBS with 10% serum, and a further 30-min incubation. After extensive washing with PBS, the appropriate Alexafluor (Fisher Scientific) conjugated secondary antibody was added in PBS with 10% serum and incubated for a further 15 min. The coverslips were washed extensively in PBS and mounted in Mowiol containing DAPI to stain nuclei. Images were acquired using a Nikon A1 confocal microscope and processed using ImageJ software.

### Gaussia luciferase reporter assay

HeLa cells were transfected with plasmid pCMV-GLuc-1 and vhs-expressing plasmids. After 16 h, the medium was replaced, sampled 4 h later, and chemiluminescence measured by injection of coelenterazine (Promega # S2001) at 1 µg/ml in PBS and read on a Clariostar plate reader.

### Fluorescent in situ hybridisation (FISH) of poly(A)+ RNA

HeLa cells were grown in 2-well slide chambers (Fisher Scientific # 154461) and transfected with plasmid. Sixteen hours later, cells were fixed for 20 min in 4% PFA in PBS. Cold methanol was added to cells for 10 mins followed by 70% ethanol for 10 mins, then 1M Tris (pH 8) for 5 mins. The cells were then incubated with 1 ng/µl 5’-labelled Cy3-oligodT (Merck) in hybridization buffer (0.005% bovine serum albumin; 1 mg/ml yeast tRNA; 10% dextran sulfate (Merck # 265252); 25% deionised formamide (Merck #S4117); in 2 x SSC (Merck # S6639) at 37 °C for 2 hours before sequential washing in 4 x SSC, and 2 x SSC; 0.1% tritonX100 in 2 x SSC. Cells were mounted in Mowiol containing DAPI to stain nuclei, images acquired with a Nikon A1 inverted confocal microscope and processed using Adobe Photoshop software.

### Fluorescent in situ hybridisation (FISH) of GLuc mRNA

Cells were grown in 2-well slide chambers (Fisher Scientific) and transfected with appropriate plasmids. After 16 hours, cells were fixed for 20 min in 4% PFA, then dehydrated by sequential 5 min incubations in 50%, 70% and 100% ethanol. FISH was then carried out using Applied Cell Diagnostics (ACD) RNAscope Multiplex reagent kit (# 323100 and # 320851) according to manufacturer’s instructions. Briefly, cells were rehydrated by sequential 2 min incubations in 70%, 50% ethanol and PBS, and treated for 30 min at 37 °C with DNase, followed by 15 min at room temperature with protease. Cells were then incubated for 2 h at 40 °C with a GLuc RNAscope probe (as designed by Advanced Cell Dignostics, ACD, #508361-C2), followed by washes and amplification stages according to instructions. After incubation with the final fluorescent probe, the cells were mounted in Mowiol containing DAPI to stain nuclei, and images acquired with a Nikon A1 inverted confocal microscope.

### siRNA depletion

Silencer select siRNA duplexes (Ambion, ThermoFisher Scientific) for XRN1 (s29016), XRN2 (s22412) or RNaseL (s12065) were forward or reverse transfected with Lipofectamine 2000 in HeLa cells to a final concentration of 20 nM according to the manufacturer’s instructions. The Silencer Select negative control siRNA number 1 was used as a negative control (Ambion, ThermoFisher Scientific).

### Quantitative RT-PCR (RT-qPCR)

Total RNA was extracted from cells using RNeasy kit (Qiagen) and on-column DNase treatment. cDNA was synthesized by qScript Ultra Supermix (VWR # 733-2904) using random primers according to manufacturer’s instructions. All qRT-PCR assays were carried out in 96-well plates using Takyon NO ROX SYBR Blue qPCR MasterMix Plus (Eurogentec #UF-NSMT-B0701). GLuc was amplified using GLuc forward primer CCTACGAAGGCGACAAAGAG and reverse primer TTGTGCAGTCCACACACAGA; GFP was amplified with forward primer GAAGCAGCACGACTTCTTCAA and reverse primer AG-TCGATGCCCTTCAGCTC. Cycling was carried out in a Lightcycler (Roche), and GLuc or GFP mRNA levels determined by the Δ ΔCt method using 18s RNA as reference gene (18s F: CCAGTAAGTGCGGGTCATAAGC; 18s R: GCCTCACTAAACCATCCAATCGG).

### Metabolic Labelling of Nascent mRNA

HeLa cells were transfected with GFP or vhs-GFP expressing plasmids, incubated for 16 hours, and cells were labelled with 0.5 µM 5-EU (source) for 40 minutes. Total RNA was extracted using Qiagen RNeasy kit with on-column DNase digestion, before biotinylation using the Click-iT Nascent RNA Capture Kit (Fisher Scientific # C10365)) as per the manufacturer’s instructions. RNA was precipitated overnight and captured using streptavidin beads. cDNA was synthesised either from total RNA samples or directly from the beads using qScript Ultra Supermix, and RT-qPCR was performed as before. In situ EU labelling was carried out with Click-&-Go Plus 555 imaging kit (2B-Scientific # CCT-1317) as per the manufacturer’s instructions.

### RNA library preparation, sequencing and analysis

Following transfection, GFP-positive cells were selected using a BD FACS Aria Fusion flow cytometer. Total RNA was extracted from 2.5 x 10^5^ cells using Qiagen RNeasy reagents, with three biological replicates prepared for each condition. The quality of the RNA preparations was assessed using a Bioanalyser. Illumina RNA-Seq libraries were constructed using the KAPA mRNA Hyperprep kit (Roche # 08098123702) according to manufacturer’s instructions. Here, 500 ng of total RNA used as input for each sample was spiked with 1 µL of a 1:100 dilution of the ERCC RNA Spike-In Mix 1 (ThermoFisher, 4456740). Sequence libraries were multiplexed and run on an Illumina NextSeq 2000 (100 cycles) resulting in paired-end (2 x 50 bp) datasets. RNA sequencing was performd at UCL Genomics (RRID SCR_027010). Bioinformatic analysis of RNAseq data was carried out as described previously ^56^.

RNAseq data can be found on Array Express Biostudies with Accession numbers E-MTAB-17530 (Total RNAseq described in Figure 1) and E-MTAB-17528 (Fractionated RNAseq data described in Figure 1G to 1I). Corresponding differential gene expression results are presented in Figure 1-Source Data 1.

### Statistical analysis

All quantified experiments were conducted on at least three biological repeats and analysed using Graphpad Prism. For comparisons between two groups, an unpaired two-tailed Student’s ***t***-test was used. For comparisons involving three or more groups, a one-way analysis of variance (ANOVA) was performed, followed by an appropriate post-hoc multiple-comparisons test where indicated. Data are presented as mean ± standard error (SE). Statistical significance was defined as *p* < 0.05.

## Acknowledgements

The authors thank the Bioimaging Technical Team in the School of Biosciences, University of Surrey, for assistance with cell sorting, and UCL Genomics for RNA sequencing. We also thank Nicolas Locker, The Pirbright Institute, for advice on experimental set up. This work was funded by UKRI Biotechnology and Biotechnology Sciences Research Council (grant BB/T007923/1 to GE).

## Notes

### Competing Interest Statement

The authors have declared no competing interest.

### Summary of Updates

This version of the manuscript has been revised to include all supplemental files.

