## Supplementary material for "XRN1 couples RNA granule remodelling to global mRNA decay triggered by the activity of a viral endoribonuclease": Figure 1 - figure supplement 1

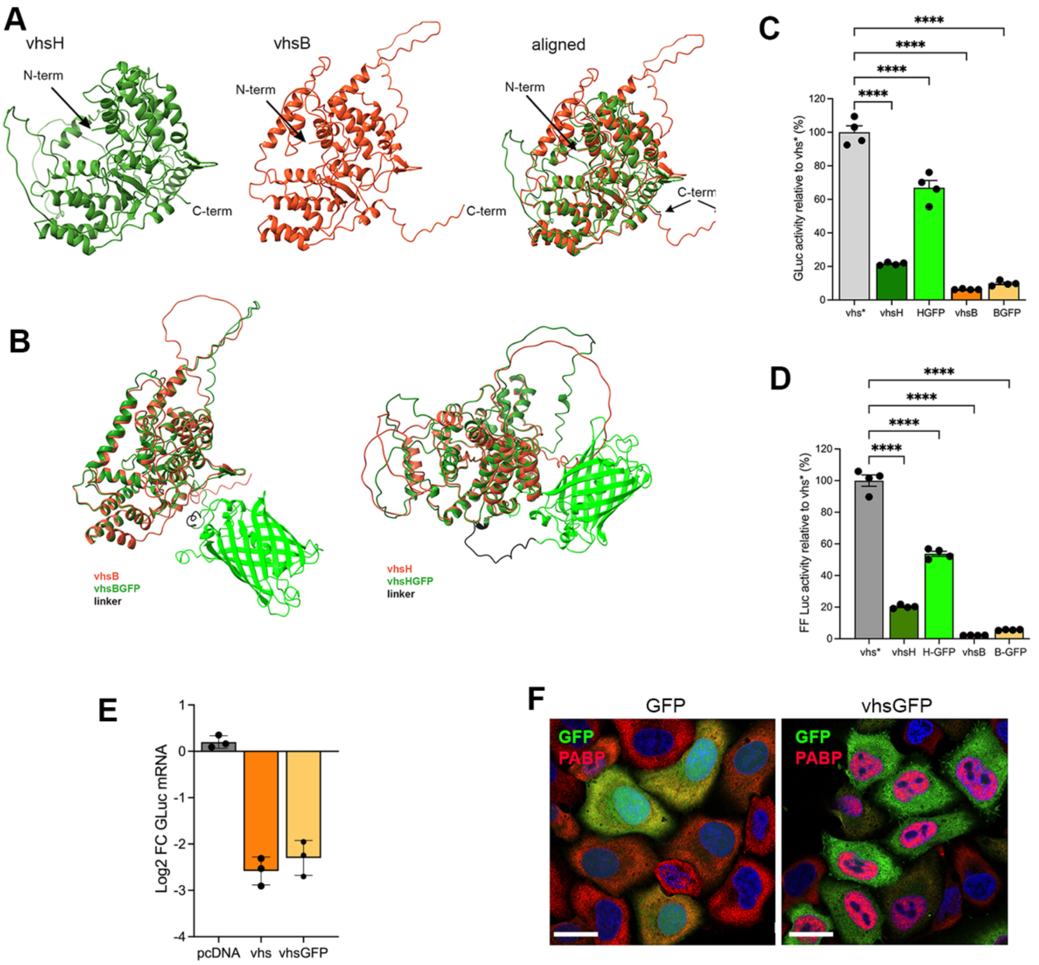


***Figure 1 – figure supplement 1.*** **(A)** Alphafold predicted structures of vhs from HSV1, vhs from BoHV1 and both aligned, indicating the N- and C-termini. **(B)** Alphafold predicted structures of vhs from HSV1 and BoHV1 fused to GFP. **(C) & (D)** HeLa cells were transfected with plasmid expressing Gaussia luciferase (C) or firefly luciferase (D) along with vhs-expressing plasmids as indicated. Relative luciferase expression was measured after 16 hours. **(E)** HeLa cells were transfected with GLuc expressing plasmid together with pcDNA, or plasmid expressing untagged or GFP-tagged vhs from BoHV1. After 16 hours, total RNA was harvested and RT-qPCR carried out for GLuc, using 18S as reference. **(F)** HeLa cells were transfected with plasmid expressing GFP or GFP-tagged vhs from BoHV1. After 16 hours, cells were fixed and stained for PABPC1 (red) and nuclei stained with DAPI (blue). Scale bar = 20 μm.
