## Supplementary material for "XRN1 couples RNA granule remodelling to global mRNA decay triggered by the activity of a viral endoribonuclease": Figure 3 - figure supplement 1

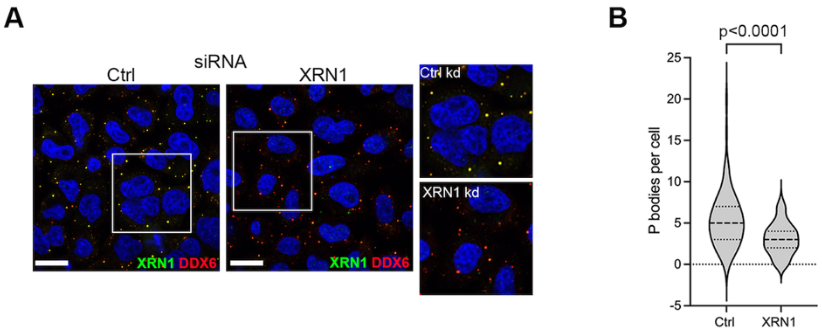


***Figure 3 –figure supplement 1.*** **(A)** HeLa cells were transfected with control or XRN1 siRNA, and after 48 hours fixed and stained for XRN1 (green) and DDX6 (red), and nuclei were stained with DAPI (blue). Scale bar = 20 µm. **(B)** P-body counts from 100 cells for each condition were carried out using ImageJ software and plotted as a violin plot showing median and quartiles. Statistical analysis was carried out using student's T-test. **** $p < 0.0001$.
