## Supplementary material for "XRN1 couples RNA granule remodelling to global mRNA decay triggered by the activity of a viral endoribonuclease": Figure 4 - figure supplement 1

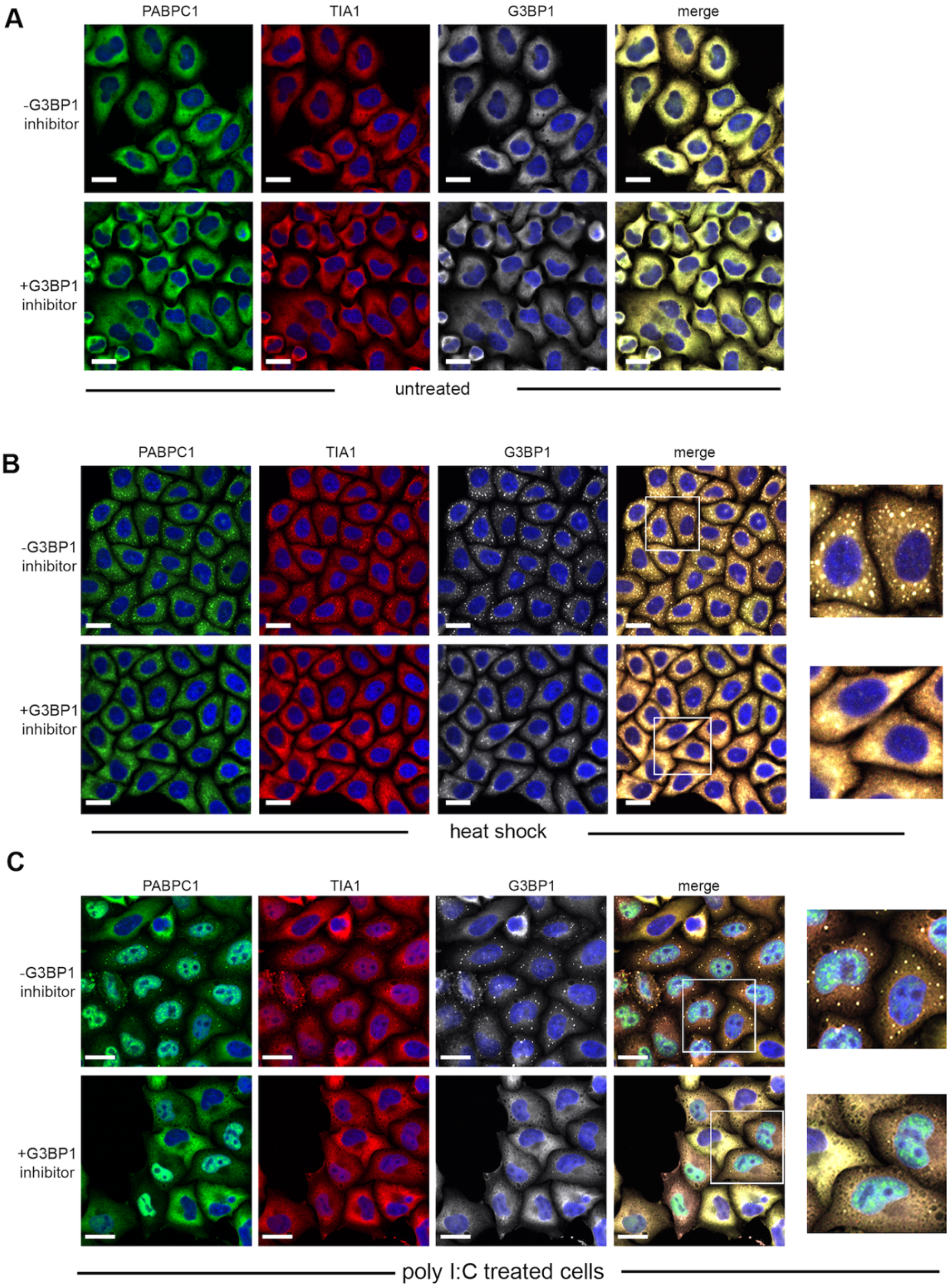


***Figure 4. supplemental figure 1.*** HeLa cells were left untreated **(A)** or subjected to heat shock at 44 °C to induce canoncial stress granules **(B)** in the absence or presence of G31b pretreatment, and fixed after 10 mins. Cells were stained for PABPC1 (green), TIA1 (red) and G3BP1 (white) and nuclei stained with DAPI. Scale bar = 20 µm. **(C)** HeLa cells were left untreated or pretreated with G31b, before transfecting poly I:C to activate RNase-L. After 3 hours, cells were fixed and stained for PABPC1 (green), TIA1 (red) and G3BP1 (white). Nuclei were stained with DAPI (blue). Scale bar = 20 mm.
