## Supplementary material for "XRN1 couples RNA granule remodelling to global mRNA decay triggered by the activity of a viral endoribonuclease": Figure 6 - figure supplement 1

D34 D83 E192 D194 D213 D215 D261 T211

```
VhsH      MGLFGMMKFAHTHHLVKRRGL-GAPAGYFTPIAVDLWNVMYTLVVKYQR--RYPSYDREA  57
VhsV      MGLFGLTRFIHEHKLVKPSII-STPPGVLTPVAVDVWNVMYTLLERLYPVGKRENLHGPS  59
VhsM      MGVYGCMNFAYANGLVRSHISGFISGGDMTPIAVDLWNVMYGLLEKLYP-GKIDGVDDSI  59
VhsB      MGLFKLLRYAYGNRLVKHDAI-TTPPGVMTPIAVDLWNVMYTLLERFCG-DAPGGVGDAA  58
VhsE      MGLFGLLKYAYSNRLVKHDAI-TTPPGIMTPIAIDLWNVMYTLMEKFDQ-ERNFPLDGAA  58
          **::   .: : : **:         * :**:*:*:***** *: :

VhsH      ITLHCLCRLLKVFTQKSLFPIFVTDRGVNCMEPVVFGAKAILARTTAQCRT---------  108
VhsV      VTIHCLGVLLRLLTQRSYYPIFVLERCT--DGPLSRGAKAIMSRAMNHDERGTSDLTRVL  117
VhsM      ITLRCLYCLLRLLHQRSYYPVFVADRGFYGNGKTLYGAKAIAATCIATRG-GSGRLHARL  118
VhsB      ATARCFLSLLRMLLKRSYYPIFVADRGIHGDRRATRGAKAIVAQTMRAVG-GSGRLGRLV  117
VhsE      VTARCFFSLLRLLLKRSYYPIFVSDRGIYGDGRVKQGAKAIVSQTMSSYG-GSGRLSSAC  117
           * :*:  **::: ::* :*:** :*          ***** :

VhsH      --DEEASDVDASP----------------------PPSPITDSRPSSAFSNMRRRGTSL  143
VhsV      LSSNTSCSIKYNKTSETYDSVFRNSSTSCIPSEENKSQDMFLDGCPRQTDKTI-------  170
VhsM      FDECGSTD-VGEA-----------------------------RC---------------  132
VhsB      SDDYTSED-EVLG---AYE-----------------YPVPHADAAADDDEEAT-------  149
VhsE      FTGDEHDT-EFQEDPEEND-----------------VSVPPQDTCPPTEISAG-------  152

VhsH      ASGTRGTAGSGAALPSAAPSKPALRLAHLFCIRVLRALGYAYINSGQLEADDACANLYHT  203
VhsV      --CLRDQNVCSLTSTMPSRGHPNHRLYHKLCASLIRWMGYAYVEAVDIEADEACANLFHT  228
VhsM      -----KRKHPGEKVCRWSERFETPKILYQLCMSIIRYMGYPYVDAGTMEADDICANLFHT  187
VhsB      ------AKEFAGRASAGAARANAPKLAHRVCVSLIRFLGYAYVDAAEMEADDVCANLFHT  203
VhsE      --YVEPERKCEHSSTRWSALDGAPRLSYRLCVNLIRHLGYPYVNACNLEADDVCANLYHT  210
                              :      :: : .*  ::* :** *:::  :***: ****:**

VhsH      NTVAYVYTTDTDLLLMGCDIVLDISACYIPTINCRDILKYFKMSYPQFLALFVRCHTDLH  263
VhsV      RTVALVYTTDTDLLFMGCDILLDAIPMFAPVVRCRDLLQYLGITYPEFLVAFVRCQTDLH  288
VhsM      KTVAYVLSSDTDLILMGCDIIIDLTRIFPPTIYCRDVLAALQMDYFTFLLNFVRCHTDLH  247
VhsB      NTVAHIYTTDTDMILMGCDLILDAAPLFPPTLRCRDVLASLGLTYGQFLATFVRCHTDLH  263
VhsE      NTVAQIYTTDTDLILMGCDIILDIMPLFPPTLRCCDVLMDLGVTYDEFLTEFVRCHTDLH  270
          .*** : ::***::****::*    : *.: * *:*  : : *  **  ****:****
```

***Figure 6 – figure supplement 1.*** Homologues of vhs from five alphaherpesviruses were aligned using BLAST. Important residues for catalysis are highlighted according to [43]: blue – Mg binding site 1; green – Mg binding site 2; turquoise - threonine residue T211 thought to be important for activity and hydrogen bonds with D34 in Mg binding site 1.
