## Supplementary material for "XRN1 couples RNA granule remodelling to global mRNA decay triggered by the activity of a viral endoribonuclease": Figure 6 - figure supplement 2

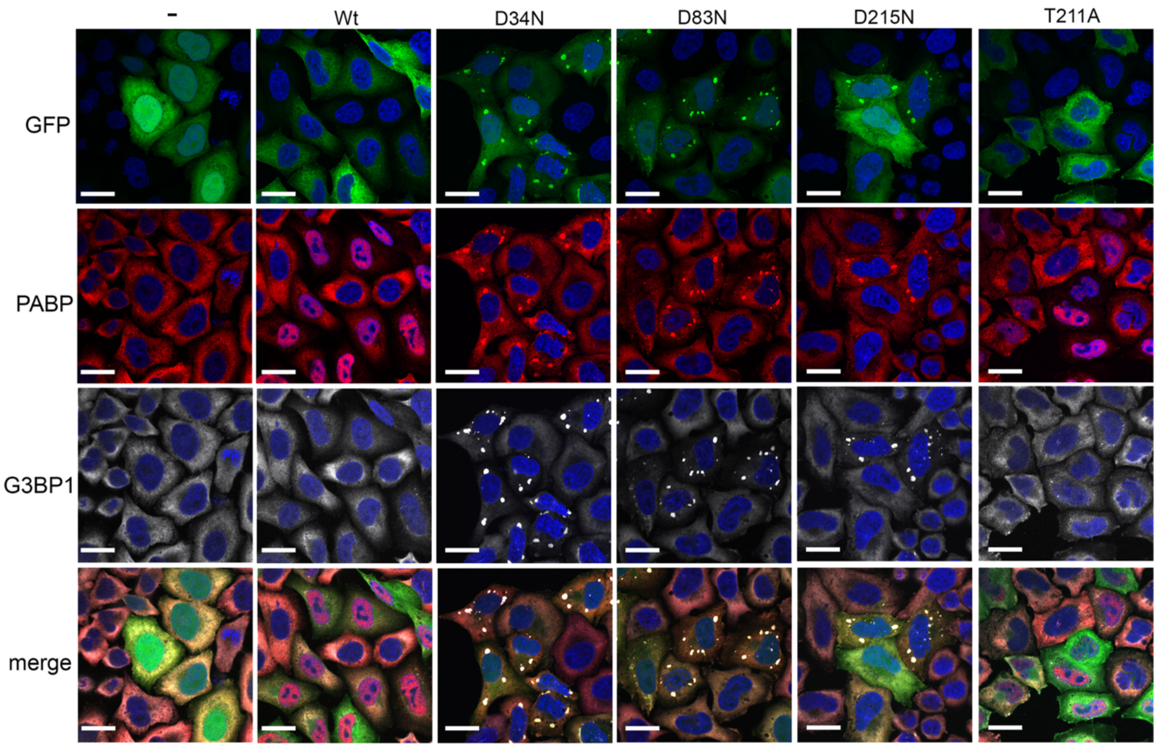


***Figure 6 – figure supplement 2.*** HeLa cells transfected with plasmids expressing GFP or GFP-tagged variants of vhs (green) were fixed and stained 16 hours later for PABPC1 (red) and G3BP1 (white). Nuclei are stained wih DAPI (blue). Scale bar = 20 μm.
