## Supplementary material for "XRN1 couples RNA granule remodelling to global mRNA decay triggered by the activity of a viral endoribonuclease": Figure 6- figure supplement 3

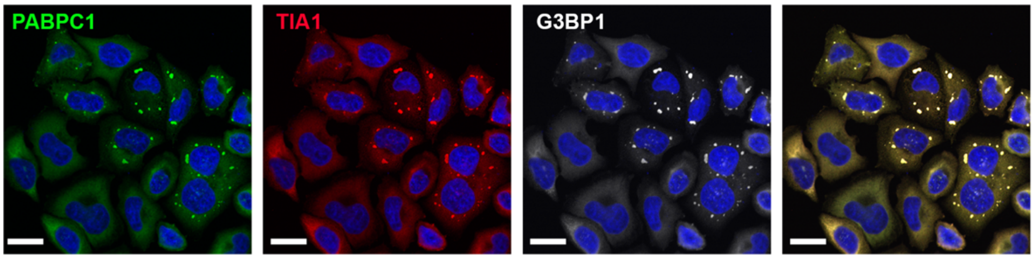


***Figure 6 – figure supplement 3.*** HeLa cells were left untreated or pretreated with G31b, before transfecting poly I:C to activate RNase-L. After 3 hours, cells were fixed and stained for PABPC1 (green), TIA1 (red) and G3BP1 (white). Nuclei were stained with DAPI (blue). Scale bar = 20 μm.
