## Supplementary material for "XRN1 couples RNA granule remodelling to global mRNA decay triggered by the activity of a viral endoribonuclease": Figure 7 - figure supplement 1

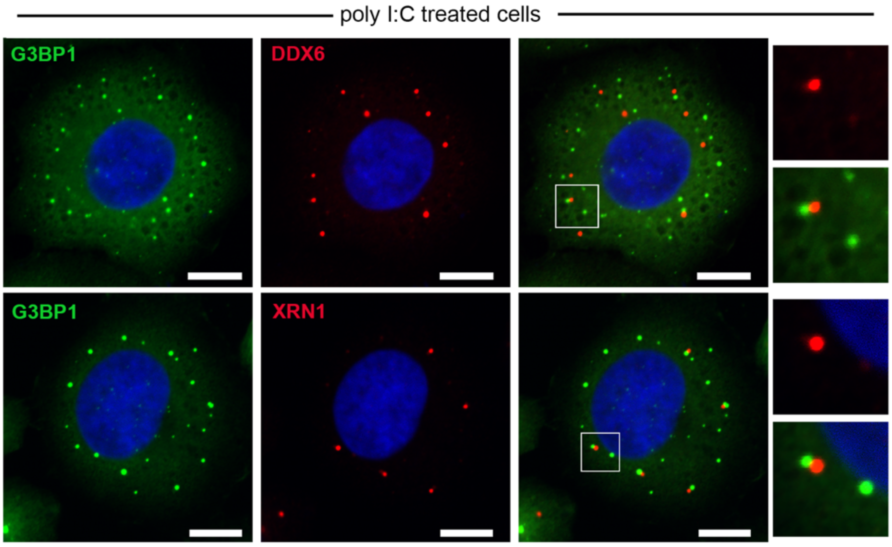


*Figure 7 – figure supplement 1.* HeLa cells were transfected with poly(I:C) to activate RNase-L. After 3 hours, cells were fixed and stained for G3BP1 (green) and DDX6 or XRN1 (red). Nuclei were stained with DAPI (blue). Scale bar = 10 μm.
